# Chondroitin and dermatan sulfate exposure induces a wound healing state in fibroblasts through Cux1-mediated SerpinB2 transcriptional repression

**DOI:** 10.1101/2023.08.01.551410

**Authors:** Alba Diaz-Pizarro, Nuria Del Valle-Del Pino, Enrique Galán, Jose María Carvajal-González, Ángel-Carlos Román, Sonia Mulero-Navarro

## Abstract

Mucopolysaccharidoses (MPS) are a group of syndromes characterized by the accumulation of sulfated glycosaminoglycans (sGAGs), leading to profound connective tissue alterations, including impaired endochondral ossification. The function of sGAGs involves determining the mechanical properties of the extracellular matrix and regulating growth factor signaling pathways, such as Fgf2. In this study, we investigated the deposition of chondroitin sulfate and dermatan sulfate, two major sGAGs, and their resemblance to wound healing states in human fibroblasts. Our findings indicate that this condition alters cell adhesion, providing a potential explanation for fibrosis-like changes observed in MPS patients. Furthermore, we elucidate the molecular pathway responsible for this effect, wherein increased Cathepsin L activation leads to the processing of the transcription factor Cux1 into a stable form capable of regulating the expression of target genes, including SERPINB2. The presence of similar changes in cell adhesion in human-induced pluripotent stem cell-derived mesenchymal cells further reinforces the significance of sGAGs in cell adhesion and sheds light on possible mechanisms underlying altered endochondral ossification in MPS patients.

## BACKGROUND

Sulfated glycosaminoglycans (sGAGs) are complex linear polysaccharides whose biological function primarily arises from their covalent attachment to core proteins, forming proteoglycans (Iozzo & Schaefer, 2015). These sGAGs can be categorized based on their sugar composition, including keratan sulfate (KS), chondroitin sulfate (CS), dermatan sulfate (DS), and heparan sulfate (HS). The disaccharide units of these polysaccharides typically consist of a uronic acid and a hexosamine, except for KS, which contains galactose instead of a uronic acid (Lamberg & Stoolmiller, 1974). Due to the addition of sulfate groups along with carboxylate groups of sugars, these molecules carry a polyanionic nature, facilitating ionic interactions between sGAGs and proteoglycans as well as other proteins (Vallet et al., 2022). A well-known biological function of sGAGs within proteoglycans is their ability to interact with various proteins in the extracellular matrix (ECM) and cell membrane. This interaction with ligands can regulate protein activity, protect ligands from degradation, or serve as a reservoir for future mobilization (Varki et al., 1999). While some interactions are common to most or all sGAGs, others are specific to particular types. Recent proteomic analyses have emphasized the importance of sulfation patterns in determining protein-sGAG interactions (Vallet et al., 2022). Additionally, a molecule called surfen (bis-2-methyl-4-amino-quinolyl-6-carbamide) has been found to interact with sGAGs, blocking Fgf2 activation in different cell types (Chatterjee et al., 2019; Huang et al., 2018; Schuksz et al., 2008).

In pathophysiology, enzymes responsible for producing, modifying, or degrading sGAGs play a critical role since their alteration can lead to genetic rare diseases known as mucopolysaccharidoses (MPSs). These diseases arise due to the deficiency of specific lysosomal enzymes responsible for degrading sGAGs. Consequently, the accumulation of partially degraded sGAGs within cells impairs cell function (Wraith, 1995). Currently, the only available treatments for MPSs are enzyme replacement therapies (ERTs). As sGAGs are present in most tissues, these diseases result in multiple organ failures and reduced life expectancy (Muenzer, 2011). Common phenotypes in MPSs include skeletal abnormalities (dysostosis multiplex, (Hampe et al., 2020)), characterized by short stature, restricted mobility, joint stiffening, coarse facial features, impaired pulmonary function, cardiac defects, and hepatomegaly, among others (Muenzer, 2011). Each MPS type exhibits distinct phenotypes as the specific sGAG accumulated varies depending on the affected enzyme. Although the ultimate consequences of MPSs are well known, the early alterations caused by sGAGs accumulation remain unclear. Recent studies have shed light on how sGAGs accumulation may impair relevant proteoglycan functions in cell signaling across tissues (Costa et al., 2017; Cyske et al., 2022; De Pasquale & Pavone, 2019; Gaffke, Pierzynowska, Podlacha, et al., 2020; Heppner et al., 2015). Therefore, various transcriptional programs responsible for different MPSs have been elucidated (Brokowska et al., 2022; Gaffke et al., 2022; Gaffke, Pierzynowska, Krzelowska, et al., 2020; Gaffke, Pierzynowska, Podlacha, et al., 2020; Wiśniewska et al., 2022). However, the transcription factors governing these changes remain unknown.

In this study, we unveil a molecular pathway by which sGAGs, specifically CS and DS, regulate gene expression in human fibroblasts. Notably, sGAGs and proteoglycans are integral components of the extracellular matrix (ECM), and fibroblasts, being major connective tissue cell types, play a crucial role in ECM function (Junqueira & Montes, 1983). Thus, alterations in sGAG composition could affect several properties of the ECM and connective tissue (Paganini et al., 2019; Sodhi & Panitch, 2021). Fibroblasts are particularly involved in wound healing, as well as its pathological counterpart, fibrosis. Regular wound healing involves the activation of fibroblast progenitors into myofibroblasts, leading to migration, invasion, and increased ECM production (Henderson et al., 2020). This process is regulated by various chemokines and inflammatory mediators released during the coagulation of blood (serum) in response to tissue injury (Iyer et al., 1999). Fibrosis represents an abnormal wound healing process, characterized by excessive scar formation and loss of tissue function (Lurje et al., 2023). While primary fibrotic diseases have low incidence, fibrosis frequently exacerbates other chronic diseases such as hepatitis or ischemia and accounts for approximately 45% of deaths in industrialized countries (Henderson et al., 2020). Although clinical observations have shown fibrosis in the liver and heart of MPS patients (Keller et al., 1987; Parfrey & Hutchins, 1986), direct mechanisms related to sGAGs have not been fully described. In this paper, we demonstrate how exposure to sGAGs induces a wound healing-like state in fibroblasts from both transcriptional and physiological perspectives. We describe a molecular mechanism involving the activation of the transcription factor Cux1, attributed to increased Cathepsin L activity, leading to a transcriptional program that impacts cellular adhesion, with SERPINB2 identified as a direct target of this pathway.

## RESULTS AND DISCUSSION

### CS/DS exposure induces a molecular response similar to serum exposure in fibroblasts

First, we aimed to characterize the transcriptional response to sGAGs exposure using a patient with Maroteaux-Lamy Syndrome (MPS VI) as a model. This patient had a congenital homozygous defect in ARSB, the gene responsible for encoding the Arylsulfatase B enzyme, which is involved in the degradation of CS and DS (Supplementary figure 1). Consequently, the patient exhibited increased sGAG levels in the urine, along with clinical phenotypes such as dysostosis multiplex, hepatomegaly, and coarse facial features. To understand the transcriptomic changes induced by sGAG accumulation, we compared the transcriptomes of primary dermal fibroblasts from the MPS VI patient (pML) with those of healthy dermal fibroblasts (pDF) (Figures 1A and 1B). Additionally, we treated pDF with CS and DS (at concentrations of 150 ug/ml and 50 ug/ml, respectively, for 48 hours; Figure 1A) to account for potential off-target effects due to genetic variations. Interestingly, the transcriptional changes observed in pML and in response to DS/CS treatment correlated significantly (R2 = 0.6, P<0.001; Figure 1C). Therefore, we performed Gene Set Enrichment Analysis (GSEA) (Subramanian et al., 2005) to identify relevant expression signatures that were statistically similar to the response induced by CS/DS treatment (Figure 1D). Several of these identified signatures were associated with extracellular matrix and collagen organization, which was expected considering the known interactions between glycosaminoglycans and collagen (Munakata et al., 1999), potentially leading to a transcriptional response. Notably, a specific signature, the fibroblast core serum response (CSR) signature, appeared in the top two GSEA pathways (Figure 1D). Genes upregulated in human fibroblasts after 36h serum exposure were also significantly upregulated in pML fibroblasts or after CS/DS exposure, while genes downregulated after 36h serum exposure were repressed in pML fibroblasts or after CS/DS exposure. The CSR signature is associated with the wound healing response of fibroblasts (Chang et al., 2004; Iyer et al., 1999), suggesting that fibroblasts activate this response when encountering serum during tissue injury. The CSR signature includes genes related to cell proliferation, migration, adhesion, and sterol biosynthesis (due to the presence of sterols in serum) (Iyer et al., 1999). To investigate if these pathways were involved in the CS/DS response, we used GSEA scores to categorize the two gene sets into four groups: upregulated in both CSR and CS/DS/ML, upregulated only in CSR, downregulated in both CSR and CS/DS/ML, and downregulated only in CSR. Our analysis revealed that cell cycle, cell migration, and immune response genes were enriched only in the gene set that was either both up- or down-regulated in CSR and CS/DS/ML. Extracellular genes were enriched in the set of genes upregulated only in CSR and in both sets downregulated (CSR and CS/DS/ML), while sterol biosynthesis genes were found only in the set downregulated in CSR (Figure 1E). These results suggest that the majority of the pathways activated during wound healing are also modulated after CS/DS exposure, even in the presence of serum, except for sterol biosynthesis genes, which are not required since serum was present in all samples. Additionally, CS is the main GAG present in blood (Bratulic et al., 2022; Gatto et al., 2016). To confirm that the fibroblast wound healing response to serum exposure was due to the presence of CS/DS, we repeated the core serum response experiment in normal fibroblasts (Iyer et al., 1999) but pre-treated the serum with chondroitinase ABC for 16 hours to remove CS/DS. We observed that the transcriptional response of ITGA6 to serum exposure was diminished in samples with CS/DS removal (Figure 1F), although we did not observe a clear effect on NTN4, another target gene of the core serum response gene set (Figure 1G). Further experiments are needed to clarify the role of CS/DS and specific genes in the fibroblast serum response.

**FIGURE 1.**
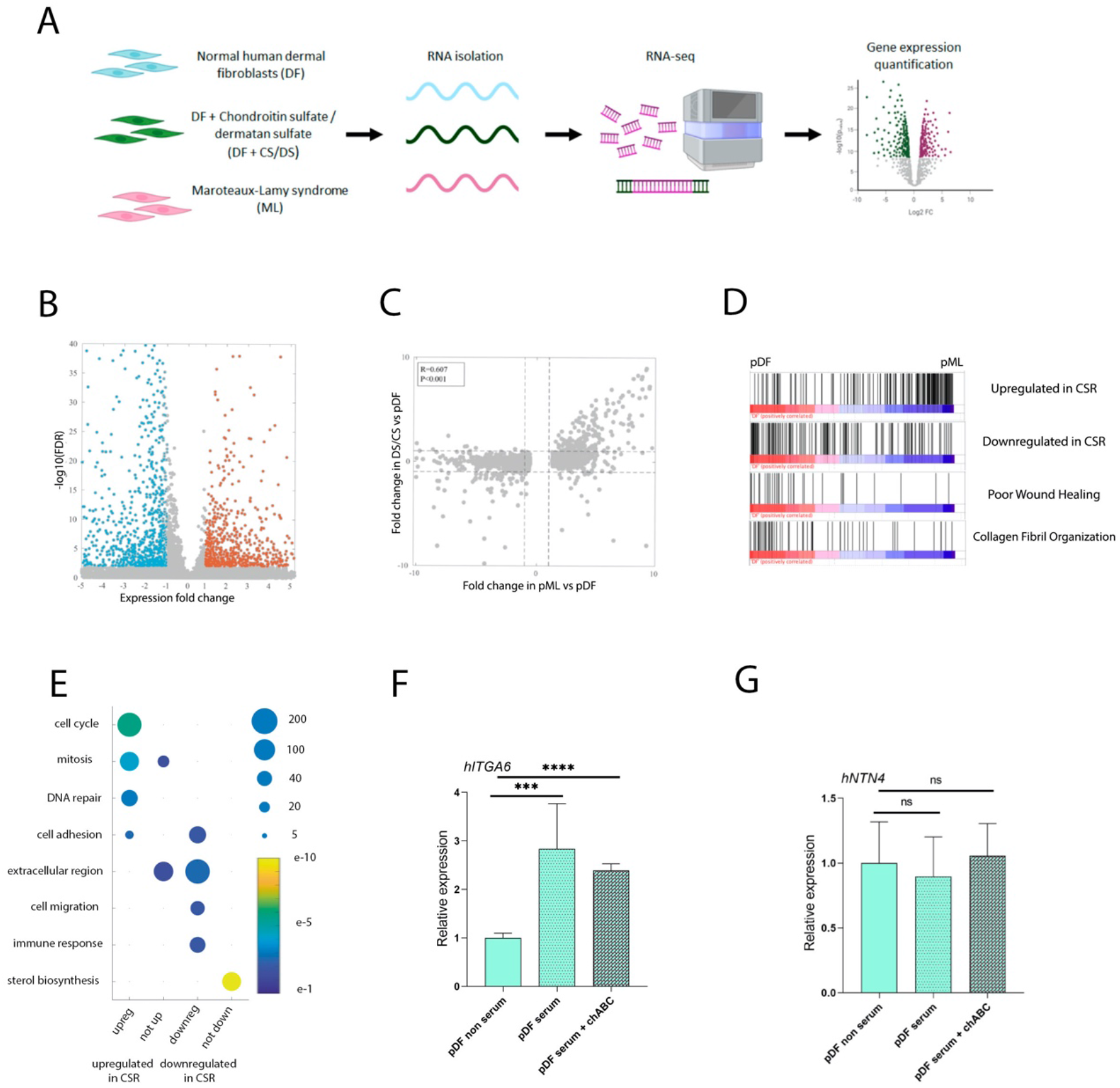
(A) Workflow of the RNAseq analysis used for dermal fibroblasts. (B) Volcano plot from the RNA-seq in primary dermal fibroblasts from the ML patient (pML) compared to primary healthy dermal fibroblasts (pDF). (C) Scatter plot for comparison between DEG (x-axis) in pML and DS/CS (y-axis) treatment compared to pDF. (D) Gene Set Enrichment Analysis (GSEA) in pDF compared to pML. (E) GO pathways (y-axis) enrichment of upregulated (bottom, left) or downregulated (bottom, right) CSR genes that are also upregulated or downregulated in CS/DS/ML (x-axis). (F-G) mRNA expression levels in pDF without serum, with serum and with serum + chondroitinase ABC (chABC). hITGA6 (F) and hNTN4 (G). Mean expression is normalized to control cells and standard deviation as error bars were plotted, n=6. P-values were obtained using two-tailed t-test (**** represents p<0.0001, *** represents p<0.001 and ns means no significative differences).

### Migration and adhesion are activated in response to CS/DS in human fibroblasts

Afterwards we sought to determine whether the molecular changes observed in pML fibroblasts and healthy pDF after CS/DS exposure were indicative of a physiological wound healing response, similar to the core serum response proposed by Iyer et al. (1999). To address this, we generated immortalized dermal fibroblast cell lines from primary fibroblasts obtained from healthy (DF) and MPS VI (ML) patients using mouse telomerase (Supplementary Figure 2). The ML clones exhibited increased secretion of GAGs into the medium, indicating their resemblance to the parental lines (Figure 2A). Additionally, we examined the gene expression of several targets that were found to be up- and down-regulated in the primary fibroblasts (Figure 1). We observed that the different clones maintained similar expression changes to those of the original primary fibroblasts (Figure 2B-F).

**FIGURE 2.**
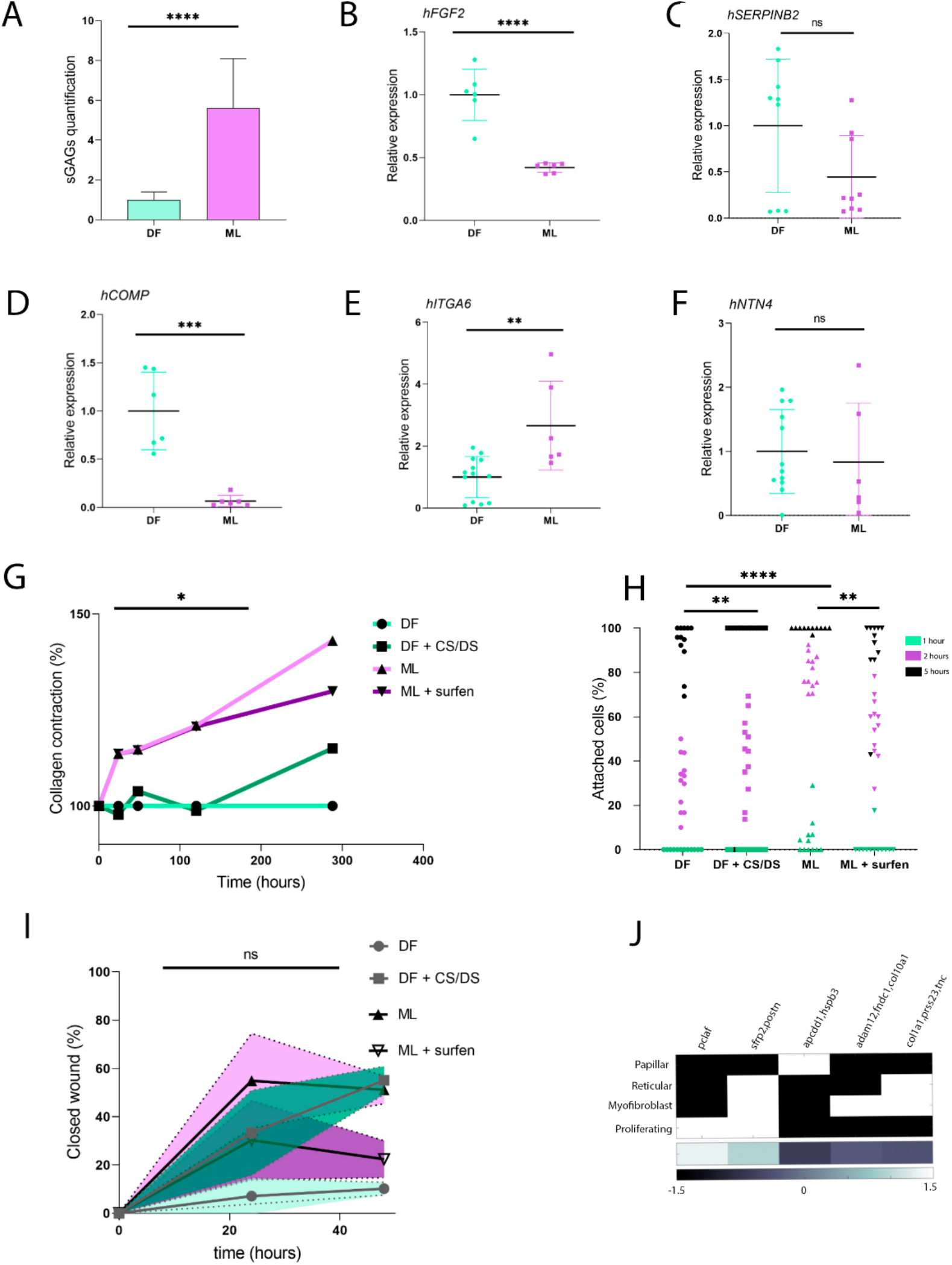
(A) Quantification of glycosaminoglycans (sGAGs) in DF vs ML using DMMB. (B-F) mRNA expression levels in DF vs ML. hFGF2 (B), hSERPINB2 (C), hCOMP (D), hITGA6 (E) and hNTN4 (F). Mean expression is relative to DF and standard deviation as error bars were plotted, n=6. (G) Quantification of collagen contraction (%) in DF with and without DS/CS treatment and ML with and without surfen treatment, n=3. * represents DF vs ML. The rest of the groups are ns. (H) Quantification of attached cells (%) in DF with and without DS/CS treatment and ML with and without surfen treatment, n=12. (I) Quantification of closed wound (%) in the migration assay in DF with and without DS/CS treatment and ML with and without surfen treatment, n=4. P values were obtained using two-tailed t-test (**** represents p<0,0001, *** represents p<0,001, ** represents p<0,01, * represents p<0,05 and ns means no significative differences). (J) Comparison of marker expression (in columns) in papillar, reticular, myofibroblastic and proliferating fibroblasts (in rows) (Tabib et al., 2021), using white/black as expressed/non-expresed markers, respectively. In the bottom row, the fold change between CS/DS/pML and pDF is represented quantitatively from black to white.

Subsequently, we compared the wound healing abilities of DF and ML cells using the collagen contraction assay (Carlson & Longaker, 2004; Zhang et al., 2022). Surprisingly, we found that the contraction produced by ML cells was lower than that of DF cells (Figure 2G). Furthermore, we discovered that this process was mediated by sGAGs, as CS/DS treatment in DF reduced the contraction, while Surfen, an antagonist of sGAGs, increased the contraction in ML-populated gels (Figure 2G). Nevertheless, we found that ML cells exhibited significantly increased cell adhesion to vitronectin compared to DF cells (Figure 2H). As seen in the collagen contraction assay, the addition of CS/DS to DF and Surfen to ML reverted the cell adhesion phenotypes (Figure 2H). Moreover, a scratch-wound assay revealed accelerated migration in ML compared to DF, with CS/DS and Surfen causing increased and decreased migration in DF and ML, respectively (Figure 2I).

To explore potential explanations for the contradictory wound healing properties of sGAG exposure in fibroblasts, we investigated the contractility of the extracellular matrix during wound healing, which is known to be regulated by myofibroblasts (LeBleu & Neilson, 2020). Myofibroblasts are activated fibroblasts characterized by the presence of αSMA (alpha smooth muscle actin) and other markers (Tomasek et al., 2002). When we compared the transcriptional profiles of DF, DF+CS/DS, and ML with a recently published signature of different types of fibroblasts (Tabib et al., 2021), we observed that the transcriptional changes induced by sGAGs did not correspond to myofibroblast activation but instead resembled a minor subset of fibroblasts exhibiting more proliferative activity (Figure 2J). Consequently, CS/DS exposure would increase cell migration and adhesion in human fibroblasts while reducing their contractility.

### Cux1 is activated in ML patients and after CS/DS exposure

Next, our focus shifted towards identifying the transcription factors responsible for orchestrating the transcriptional response to CS/DS, leading to the altered wound healing phenotype. Specifically, we analyzed the enrichment of transcription factor binding sites in the promoters of genes (Nassar et al., 2023) that exhibited significant changes in expression either in ML samples or after CS/DS exposure (Figure 1A). Notably, we found that Cux1 was the top transcription factor enriched in both CS/DS-treated cells and ML cells (Figure 3A).

**FIGURE 3.**
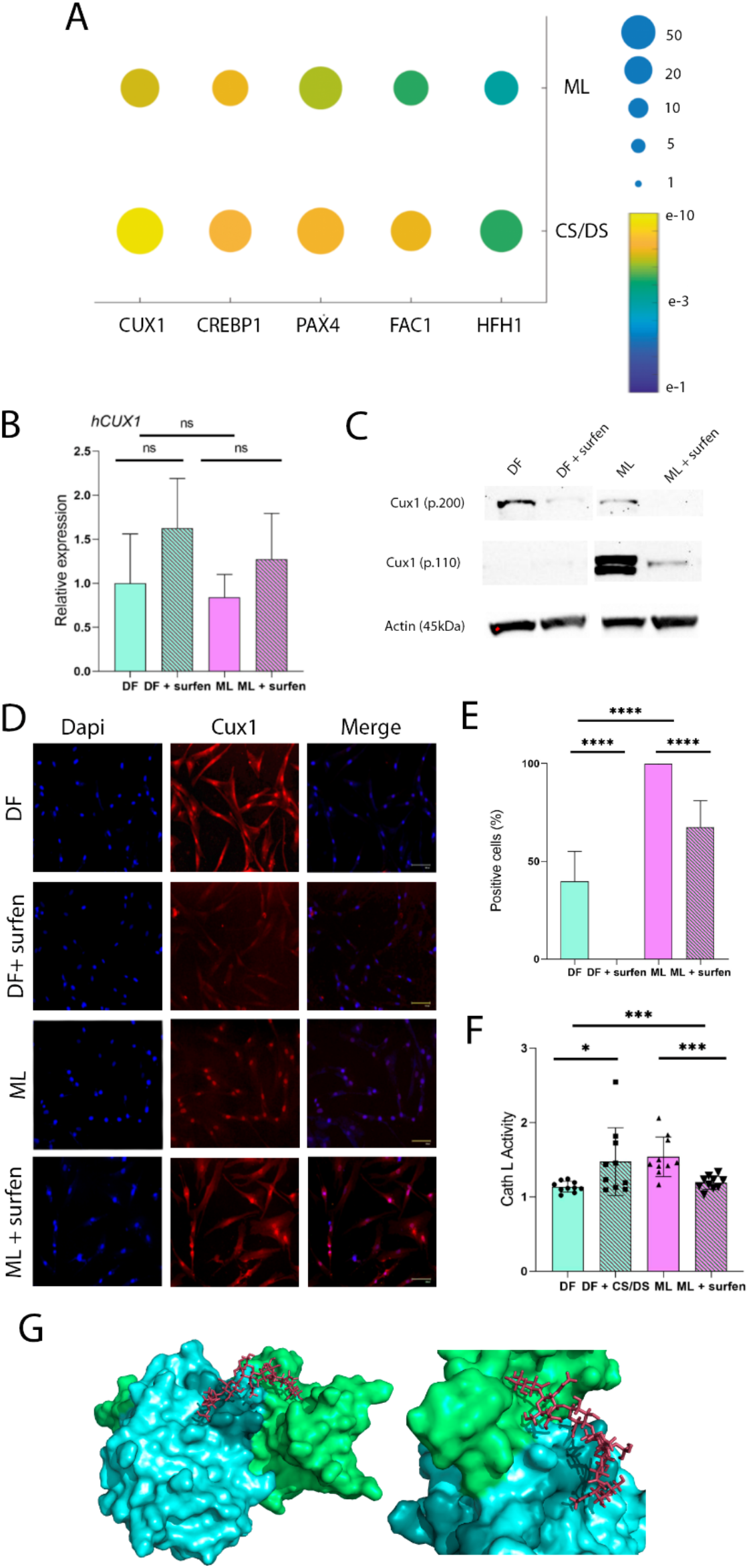
(A) Enrichment of transcription factor binding sites in the promoters of the genes in CS/DS/pML vs pDF. (B) mRNA expression levels in DF vs ML with and without surfen treatment in hCUX1. Mean relative to control cells and standard deviation as error bars were plotted, n=3. (C) Western blot for Cux1 and ß-actin in DF and ML with and without surfen treatment. (D) Immunofluorescence images for Dapi and Cux1 in DF and ML with and without surfen treatment. (E) Quantification of positive cells (%) from images in D. (F) Quantification of Cathepsin L activity in DF with and without CS/DS treatment and ML with and without surfen treatment. (G) Procathepsin L docking simulation against CS. Green and blue regions in procathepsin L represent processed and mature regions, respectively. P-values were obtained using two-tailed t-test (**** represents p<0,0001, *** represents p<0,001, * represents p<0,05 and ns means no significative differences).

Cux1 is a protein found in all metazoans, with potential activation and repression abilities, and it exists in different isoforms. It has been implicated in various physiological processes such as cell proliferation, migration, and motility (Liu et al., 2020; Nepveu, 2001; Sansregret & Nepveu, 2008), making it a compelling candidate for the regulation of the GAGs response. We did not observe significant changes in CUX1 mRNA levels in DF or ML cells (Figure 3B); however, at the protein level, we detected significant differences (Figure 3C). Specifically, the Cux1 p110 isoform was increased in ML samples, while the p200 isoform was reduced. Moreover, Surfen, the GAGs antagonist, decreased both p110 and p200 isoforms (Figure 3C). The p110 isoform has been suggested to be the transcriptionally activated form of Cux1 (Vadnais et al., 2013), leading us to hypothesize that Cux1 was more active in ML cells. Supporting this notion, immunofluorescence analysis revealed that Cux1 was preferentially located in the nuclei of ML cells, whereas the Cux1 signal was lower and more diffuse in DF cells (Figure 3D, quantified in 3E). Surfen exposure to either DF or ML cells led to the removal of this signal from the nuclei (Figure 3D-E).

The p110 isoform is produced through the proteolytic processing of the full-length p200 Cux1 by cathepsin L (Goulet et al., 2004). We observed that cathepsin L activity was increased in ML compared to DF cells, while Surfen significantly reduced this activity (Figure 3F). Therefore, a possible explanation for Cux1 activation in ML cells might be that CS/DS activated cathepsin L, leading to the proteolytic processing of Cux1. Pro-cathepsin L consists of an inhibitory region and the zymogen, and docking simulations predict that CS interacts with the interface between these two regions (Figure 3G). Supporting this idea, CS/DS treatment in DF cells increased cathepsin L activity (Figure 3F). Furthermore, sGAGs have been traditionally described as potential activators of cathepsin L (Mason & Massey, 1992), suggesting that exposure to CS/DS directly activates cathepsin L, leading to the processing of Cux1 into its transcriptionally activated form.

### Cux1 and HDACs the transcription of genes related to cell adhesion

Having observed the potential activation of Cux1 after sGAG accumulation, we aimed to investigate how Cux1 regulates transcriptional responses in fibroblasts. To achieve this, we analyzed Cux1 genome binding in ML fibroblasts through chIP-seq, using Surfen treatment as a control for Cux1 inactivation, as previously observed (Figure 3). As expected, inhibition of sGAGs with Surfen resulted in a global decrease in Cux1 binding to the genome (Figure 4A). Cux1 peaks in ML were distributed throughout the genome, located in promoters, distal sites, and introns (Figure 4B). The pattern of Cux1 binding resembled findings from previous studies in other cell types, where Cux1 was found both proximal and distal to the transcriptional start sites (TSSs) of target genes (Arthur et al., 2017).

**FIGURE 4.**
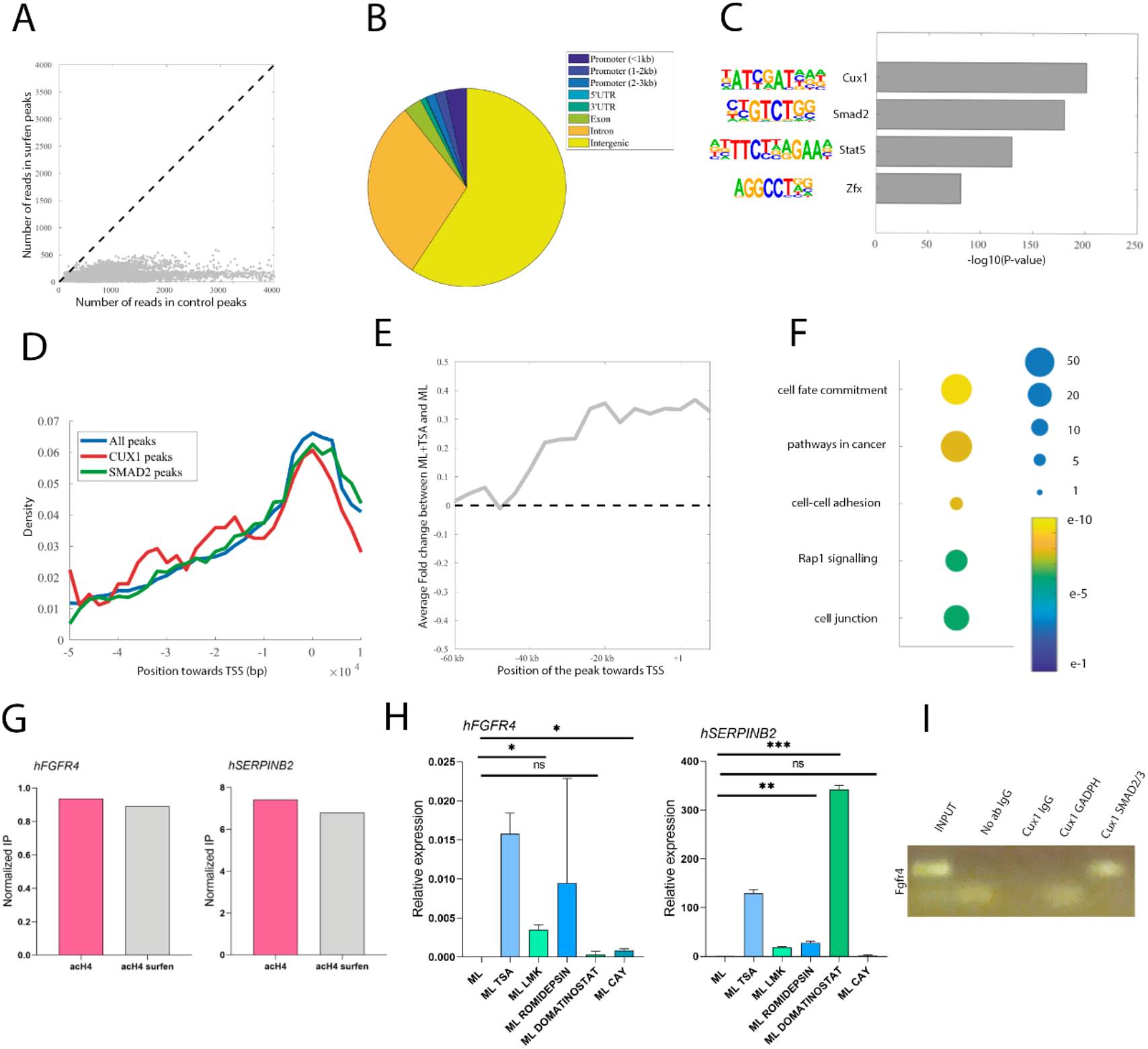
(A) Dot plot representing chIP peaks (read depth) for Cux1 in ML cells (x-axis) compared to ML cells treated with surfen (B) Distribution of Cux1 peaks in ML cells in different types of genomic regions. (C) Motif discovery in Cux1 regions from chIP-seq, including P-values from this detection (D) Position of Cux1 peaks towards TSS (from 50kb upstream to 10kb downstream). In colors, we include all peaks (blue) or peaks with associated Cux1 motif (red) or Smad2 motif (green). (E) Average fold change expression in TSA-treated ML cells compared to untreated cells in genes with Cux1 peaks present in regions near their TSS (from -60kb to +10kb). (F) Gene ontology enrichment of genes with Cux1 peaks near their TSS and significative upregulated response after TSA treatment (G) Quantification of Acetyl H4 levels in ML cells with and without surfen treatment from chIP in hFGFR4 and hSERPINB2. (H) mRNA expression levels in ML cells with different Histone Deacetylase (HDAC) inhibitors treatments; hFGFR4 (left) and hSERPINB2 (right). Mean expression is relative to untreated cells and standard deviation as error bars were plotted, n=3. (C) reChIP with either Cux1 and GADPH or Cux1 and Smad2/3 antibodies in the FGFR4 region, using input and igG as controls. P-values were obtained using two-tailed t-test (*** represents p<0,001, ** represents p<0,01, * represents p<0,05 and ns means no significative differences).

Notably, motif analyses of these peaks revealed not only enrichment of the Cux1 motif but also motifs for other transcription factors, including Smad2, Stat5, and Zfx (Figure 4C). Proximal peaks of Cux1 binding were preferentially located near the TSS, and peaks with annotated Cux1 or Smad2 motifs exhibited similar behavior (Figure 4D). Both Cux1 and Smad2/3 have been reported as partners of histone deacetylases (HDACs) (Kim & Lassar, 2003; Li et al., 1999). To explore the transcriptomic response to trichostatin A (TSA), a class I/II HDAC inhibitor (HDACi), in ML cells, we focused on genes with Cux1 peaks located near their TSS (Figure 4E). We observed that Cux1-bound regions near TSSs showed consistent correlation with genes that were overexpressed after TSA treatment (Figure 4E). Ontology analysis of Cux1-bound, TSA-overexpressed genes revealed an overrepresentation of cell adhesion terms (Figure 4F).

Further investigation of selected regions (CGNL1, FGFR4, PRKCZ, SERPINB2, and SERPINB9) showed that acetyl-H4 levels were not modulated by surfen treatment when Cux1 was unbound after treatment (Figure 4G and supplementary Figure 3A). It is possible that acetyl-H4 levels are regulated in nearby regions or indirectly through other factors. Using a panel of other HDACi, mRNA levels of Cux1-target genes pointed to Hdac1/2 as the main factor responsible for the repressive effect on target genes, as Romidepsin or Domatinostat were able to revert four out of five genes, except for SERPINB9 (Figure 4H and supplementary Figure 3B).

Finally, we investigated whether Smad2/3 might also be involved in these complexes, as they are known HDAC partners (Kim & Lassar, 2003), and their motif appeared in several Cux1 peaks, such as FGFR4 or PRKCZ regions (Figure 4C). By performing reChIP with Cux1 and Smad2/3 antibodies, we observed that the FGFR4 region was bound by both Cux1 and Smad2/3 in ML cells (Figure 4I). However, we did not find co-binding in the case of the PRKCZ region (Supplementary Figure 3C). This suggests that the binding might be cooperative in some cases while competitive in others. Further experiments will be needed to clarify the role of the relationship between Cux1 and Smad2/3, in addition to Hdac1/2 activity.

### The modulation of CS/DS levels, cathepsin L activity and Cux1 levels regulate target gene expression

At this stage, we have demonstrated that CS/DS is responsible for regulating cell migration and adhesion, as well as cathepsin L activity and Cux1 transcriptional activation. Additionally, we have identified several target genes, including CGNL1, FGFR4, PRKCZ, SERPINB2, and SERPINB9, which are related to cell adhesion and are bound by Cux1. Now, our aim is to connect the pathway involving CS/DS, cathepsin L, and Cux1 with the expression of these target genes.

Firstly, we compared the expression of these target genes in ML cells with that in DF cells (Figure 5A) and observed how surfen treatment could reverse these changes (Figure 5B). We found that CGNL1 and SERPINB2 exhibited significant changes in ML cells, which could be effectively reversed using surfen. The changes in FGFR4 and PRKCZ expression were more subtle, while SERPINB9 did not respond to surfen treatment. Furthermore, inhibiting cathepsin L activity in ML cells also reversed the gene expression changes observed in SERPINB2, while CGNL1 and FGFR4 showed a tendency that was not statistically significant (Figure 5C).

**FIGURE 5.**
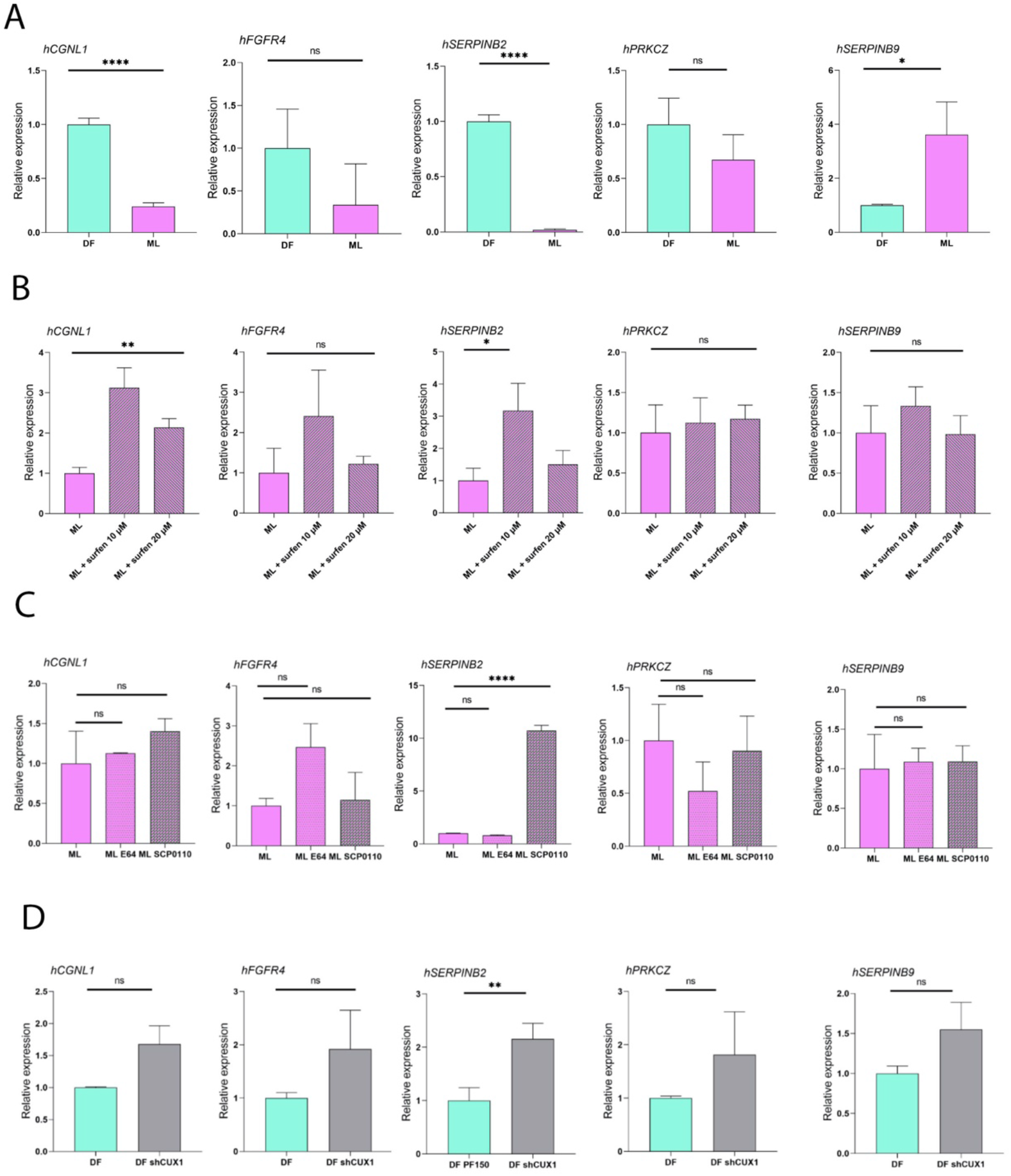
(A) mRNA expression levels in DF vs ML in hCGNL1, hFGFR4, hSERPINB2, hPRKCZ and hSERPINB9. (B) mRNA expression levels in ML with and without surfen treatment in hCGNL1, hFGFR4, hSERPINB2, hPRKCZ and hSERPINB9. (C) mRNA expression levels in ML with and without cathepsin L inhibitor treatments in hCGNL1, hFGFR4, hSERPINB2, hPRKCZ and hSERPINB9. (D) mRNA expression levels in DF vs DF shCUX1 in hCGNL1, hFGFR4, hSERPINB2, hPRKCZ and hSERPINB9. Average expression is relative to control cells and standard deviation as error bars were plotted, n=3. P-values were obtained using two-tailed t-test (**** represents p<0,0001, ** represents p<0,01, * represents p<0,05 and ns means no significative differences).

To further establish the direct role of Cux1 as a transcriptional regulator of these target genes in response to CS/DS, we generated an inducible shCUX1 knockdown fibroblast line. The reduction of Cux1 levels (Supplementary figure 4) strongly reversed the expression of the target gene SERPINB2 (Figure 5D), providing evidence that Cux1 directly regulates at least SERPINB2 by modulating its activity in response to CS/DS.

Collectively, the changes in the expression of these target genes, particularly SERPINB2, may play a crucial role in modulating the migration and adhesion properties of both ML cells and DF cells treated with CS/DS.

### Cux1 knockdown or SerpinB2 overexpression reverse cell adhesion alterations of ML cells

To assess the effect of Cux1 and its target gene SERPINB2 on the wound healing response observed in ML cells, we utilized the knockdown Cux1 fibroblast line previously generated. The wound healing response induced by CS/DS was partially restored using the knockdown Cux1 fibroblast line. Specifically, cell adhesion and collagen contraction were regulated by Cux1, while cell migration remained unaffected (Figure 5A-C). Furthermore, the overexpression of SerpinB2 was able to decrease cell adhesion in ML cells, making them insensitive to surfen treatment (Figure 5D). This suggests that the overexpression of SerpinB2 can at least partially modulate the effects of CS/DS exposure in fibroblasts. Given that alterations in cellular adhesion can explain connective tissue pathologies like fibrosis, it is important to investigate if these cellular changes are also present in other cell types, such as mesenchymal cells, which may better reflect the defects observed in MPS patients, including impaired endochondral ossification.

To address this, we generated and characterized hiPS cells from the same ML patient and differentiated them into pre-mesenchymal cells (Supplementary figure 5). We observed that ML pre-mesenchymal cells exhibited a similar increased adhesion phenotype as observed in dermal fibroblasts when compared to control episomal hIPS cells differentiated into pre-mesenchymal cells (Figure 5E). Moreover, surfen treatment reduced cell adhesion in ML pre-mesenchymal cells (Figure 5F), further confirming the similarity between these cellular models. The relevance of mesenchymal cell biology is linked to several phenotypes associated with altered sGAG levels, particularly in relation to endochondral ossification (Jiang et al., 2020). Cux1 and SerpinB2 have previously been implicated in interfering with mesenchymal to chondrocyte differentiation (Granados-Montiel et al., 2021; Lizarraga et al., 2002). Thus, our study has shed light on the role of CS/DS in cell migration and adhesion through a detailed molecular pathway (Figure 6G), which may have implications in the altered endochondral ossification observed in mucopolysaccharidosis patients.

**FIGURE 6.**
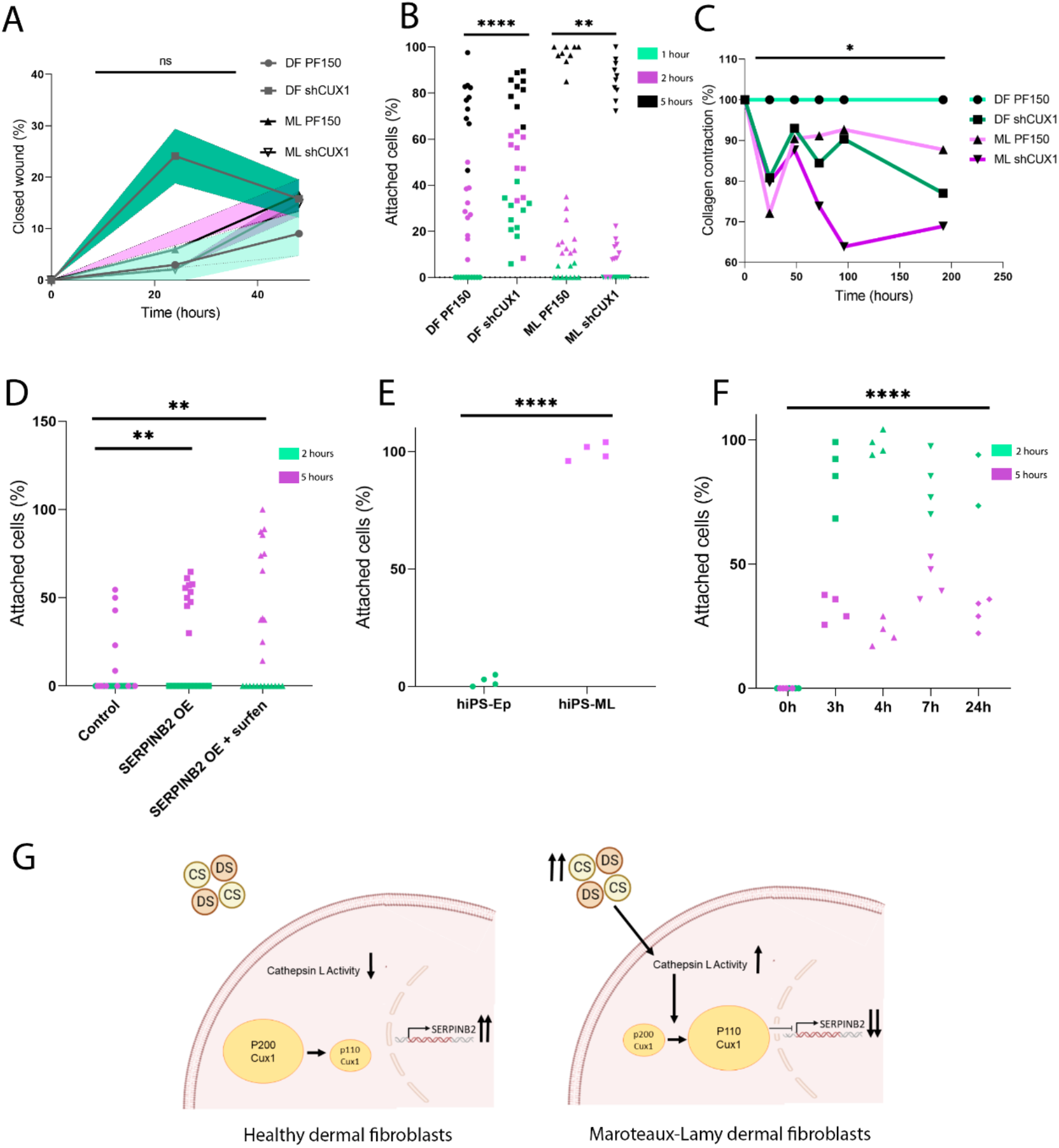
(A) Quantification of closed wound (%) in the migration assay in DF and ML and in DF shCUX1 and ML shCUX1, n=8. (B) Quantification of attached cells (%) in DF and ML and in DF shCUX1 and ML shCUX1, n=10. (C) Quantification of collagen contraction (%) in DF and ML and in DF shCUX1 and ML shCUX1. * represents DF vs ML and DF vs DF shCUX1. The rest of the groups are ns. (D) Quantification of attached cells (%) in DF and in DF with the Serpinb2 overexpressed plasmid transfected with and without surfen treatment, n=12. (E) Quantification of attached cells (%) in Control and ML in pre-mesenchymal cells, n=4. (F) Quantification of attached cells (%) ML pre-mesenchymal cells with and without surfen treatment, n=4. (G). Model of the molecular pathway that is activated in ML cells or after CS/DS treatment (right) compared to healthy fibroblasts (left). P-values were obtained using two-tailed t-test (**** represents p<0,0001, ** represents p<0,01, * represents p<0,05 and ns means no significative differences).

Future studies will aim to assess the expression and activation of Cux1 and its target genes, such as SerpinB2, in human tissues. Specifically, therapeutic treatment with SerpinB2 in patients might be relevant due to the observed deficiency in this serpin (Maas & de Maat, 2021). Additionally, the importance of fibrotic phenotypes in MPS diseases should be reevaluated, as we have demonstrated that CS/DS induces a wound healing state in fibroblasts. The role of CS/DS and other sGAGs may hold significance in potential treatments against fibrosis-associated diseases.

## MATERIALS AND METHODS

### Isolation and culture of primary human dermal fibroblasts

Human dermal fibroblasts (pML) were isolated from a skin biopsy from a patient with mucopolysaccharidosis and cultured for one month in DMEM 1x (GIBCO) supplemented with 10%. Fetal Bovine Serum (FBS) decomplemented, 5% sodium pyruvate (GIBCO) and 5% penicillin and streptomycin (GIBCO) (DMEM complete). Dermal biopsies were taken with informed consent and institutional review board approval. Different batches of human dermal fibroblasts used as control (pDF) were bought from Lonza (adult) and Zen-bio (papillary and reticular) to avoid differences from genetic or morphological backgrounds. Fibroblasts were grown in 5% CO2 at 37°C, and DMEM complete was replaced every 2 days until 70-80% confluence. At this confluence, cells were detached with 0,05% Trypsin-EDTA (1X) (GIBCO) for 5 minutes at 37°C.

### Generation of hiPSC Line and characterization

Fibroblasts were reprogrammed with CytoTune-iPS 2.0 Sendai reprogramming kit following the manufacturer’s protocol. Emerged cell clumps were picked and expanded to characterize them. The hiPS clones were expanded in VTN pre-coated six-well plates with Essential 8 Medium (Gibco) at 37°C in 5% CO2. Cells were passed with 0.02% EDTA in PBS for 3 minutes at room temperature, RevitaCell supplement (Gibco) was used to minimize apoptosis and necrosis. The pluripotency markers were analyzed by qPCR, immunofluorescence and alkaline phosphatase staining, using Alkaline Phosphatase Blue Membrane Substrate Solution (Merck) according to the manufacturer’s instructions. Silencing test for detecting the SeV (Sendai virus) genome and transgenes was performed by RT-PCR according to the protocol provided by the reprogramming kit previously mentioned.

### Pre-mesenchymal differentiation of hIPS cells

To induce differentiation, hiPS cells were dissociated with Stempro-Accutase (Gibco) and plated on Matrigel (Corning) pre-coated plates in differentiation media, which includes: DMEM with 10% Knockout Serum Replacement (Gibco), Minimum Essential Medium Non-Essential Amino Acids 1x (Gibco), L-glutamine 1x (Gibco), Pen/Strep 1x (Gibco), 2-Mercaptoethanol 1x (Gibco), bFGF (Gibco) 5 ng/ml and PDGF (RD Systems) 5 ng/ml. Cells were passed using Trypsin (Gibco) on plates with Embryomax Gelatin Solution 0,1% (Millipore) coating.

### Treatment and reagents

Fibroblasts were treated with surfen (15µM), Condroitin Sulfate (CS) (150µM) and Dermatan Sulfate (DS) (50µM). Also, they were treated with different Histone Deacetylase Inhibitors: Trichostatin A (TSA) (100 µM), LMK-235 (2 µM), Romidepsin (10nM), Domatinostat (2µM) and CAY10603 (500nM). To inhibit Cathepsin L, we used E64 (10 µM) and SCP0110 (20 µM). If unspecified, all treatments lasted 48 hours, and all reagents were bought from Sigma-Aldrich and Medchemexpress.

### Lentivirus production and infection

To generate a stable cell line, lentiviral particles were produced to drive the expression of human telomerase (hTERT) in the infected cells. pLV-hTERT-IRES-hygro (Addgene, #85140) was transfected with the packaging vectors psPAX2 (Addgene, #12260) and pMD2.G (Addgene, #12259) into HEK-293T cells using polyethylenimine (PEI; Sigma). 48 and 72 hours later, viral supernatants were concentrated in Amicon Ultra-15 filters (Merck) by centrifugation at 3500 g for 30 minutes up to an approximate concentration of 2 × 105 infectious virus particles/mL. 75 μL of concentrated virus were added in 2 mL of medium in presence of 8 μg/mL of polybrene (Sigma) for a 60-mm plate. Inmortalized fibroblasts were selected after infection with a final concentration of 100 µg/mL hygromicin during 48 hours. Short hairpin RNAs (shRNAs) against CUX1 was also generated by the production and infection of lentiviral particles using pTRIPZ-DoxOn-shCUX1-5150 (Addgene, #90469) and pTRIPZ shCUX1-5328 (Addgene, #100815) following the same protocol as explained before. For the overexpression of SERPINB2, pLENTI-GII-CMV-GFP-2A-Puro (abm, #434730610395) was transfected in fibroblasts using Lipofectamine 2000 (Invitrogen, #11668019).

### Western blot analysis

Cell lysis was carried out in ice-cold lysis buffer: 50 mM Tris-HCl (pH 7.5), 1 mM EGTA, 1 mM EDTA, 1 mM sodium orthovanadate, 5 mM sodium pyrophosphate, 10 mM sodium fluoride, 0.27 M sucrose, 0.1 mM phenylmethylsulphonyl fluoride, 0.1% (v/v) 2-mercaptoethanol, 1% (v/v) Triton X-100, and complete protease inhibitor cocktail (Roche). Bio-Rad protein assay was used to determine the protein concentration. 15 μg of protein was exposed to SDS-PAGE electrophoresis and transferred onto nitrocellulose membranes (Bio-Rad Laboratories). Those membranes were then blocked with 5% dry milk in Tris-buffered saline containing 0.05% Tween-20 for an hour and incubated overnight at 4°C with primary antibodies. The primary antibodies anti-Cux1 (Abnova, #H00001523-M01) and anti-Actin (SIGMA-ALDRICH, #A2066) were diluted in blocking solution (1:500). After several TBS-Tween washes, the membranes were incubated with horseradish peroxidase (HRP)-conjugated secondary antibodies (anti-mouse-HRP, Cell Signaling #7076; anti-rabbit-HRP, Cell Signaling #7074) in blocking solution (1:1000) for 1 hour at room temperature. After several washes with TBS-Tween, the proteins were visualized using a chemiluminescence detection system (SuperSignal West Dura, Thermo Fisher Scientific) and detected with iBright CL1000.

### RNA extraction, cDNA synthesis, qPCR, and RNA-Seq

Total RNA was isolated using RNAspin Mini RNA Isolation Kit (cytiva). After elution, 300 ng of RNA were subjected to reverse transcription using High-Capacity cDNA Reverse Transcription Kit (Applied Biosystems) according to the manufacter’s instructions. Quantitative PCR (qPCRQpcr) was performed in order to analyse gene expression using Power SYBR Green PCR Master Mix (Applied Biosystems) instructions and different oligonucleotides (the sequences are available upon request). Melt curve analyses were performed to confirm the specificity of the PCR reactions. For RNA-seq analysis, libraries from RNA samples were prepared following Illumina recommendations at Novogene UK. After that, samples were sequenced and 10M paired end reads were obtained. The reads were pseudoaligned to human genome (hg38) using kallisto, and then MATLAB was used for processing and visualization. Negative binomial distribution was required to obtain differentially expressed genes (DEGs), and DAVID or GSEA was also used for gene ontology classification.

### Cell Immunofluorescence

Fibroblasts cultures were fixed for 10 minutes at RT in 4% formaldehyde (PolySciencde) permeabilized in 0.1% Triton for 10 minutes. For 45 minutes, they were blocked with 2% bovine serum albumin (BSA, Roche). Samples were incubated overnight at 4°C in the same blocking buffer. Anti-Cux1 (Abnova, #H00001523-M01, 1:150) was the primary antibody used. Next day, samples were washed five times in 0.1% Triton and incubated for 1 hour with fluorescent secondary antibody (Alexa Fluor 568 anti-Mouse (Invitrogen, #A11005, 1:500) diluted in the same buffer. Dapi (Thermo Fisher Scientific, #62248) was used at 0.5 μg/mL to stain nuclei. Five additional washes were performed in 0.1% Triton and slides were mounted in Vectashield (Vector Labs). Images were captured using a Floid Cell Imaging Station (Thermo Fisher Scientist) and processed with ImageJ (Fiji) and Adobe Photoshop 2020. In the case of Cathepsin L, 48 hours before starting the assay, treatments were added (15 µM surfen, 150 µM Condroitin Sulfate (CS) and 50 µM Dermatan Sulfate (DS)). The measurement of Cathepsin L activity was performed following the instructions and the protocol of Magic Red Cathepsin Assays Kit (Abcam, ab270774). Photos were taken using a Floid Cell Imaging Station (Thermo Fisher Scientist), and an ad hoc MATLAB function was developed for quantification and normalization in the images.

In the case of hiPS immunofluorescence, they were first cultured on Matrigel-coated tissue culture plates (Falcon). The process was the same as in fibroblasts and the primary antibodies were anti-human NANOG (1:100, Millipore #AB5731) and SSEA4 (1:25, Developmental Studies Hybridoma Bank #MC-813-70). Secondary antibodies were Alexa Fluor 488 anti-mouse (1:500, Invitrogen #A21202) and Alexa Fluor 594 anti-goat (1:500, Invitrogen #A11058). Fluorescence was detected on the Cell Floid Imaging Station (Thermo Fisher Scientist).

### Dimethylmethylene Blue Assay (DMMB)

In order to measure the amount of glycosaminoglycans(sGAGs), a DMMB (Sigma-Aldrich) solution was prepared following manufacturer recommendations. 150,000 cells were plated per well in a 12-well plate, and after 48h in DMEM (5% sodium pyruvate, 5% penicillin and streptomycin, 0% phenol red, 0% FBS), mediums were collected. 20µL of those mediums and 200 µL of DMMB were added, and samples were measured at 525nm and quantified using control CS solutions for normalization.

### Cell migration, adhesion and contraction

Once cells were attached, treatments were added (15 µM surfen, 150 µM Condroitin Sulfate (CS) and 50 µM Dermatan Sulfate (DS)). 48 hours later, cells were detached and plated (50,000 cells/well) in untreated plastic. Photos were taken at 30 minutes, 2 hours and 5 hours using a Floid Cell Imaging Station (Thermo Fisher Scientist), and then manually counted for quantification. In the case of wound healing experiments, cells were plated (300,000 cells/well) and with a yellow tip, a wound was made at 100% confluence. Medium was changed to DMEM (5% sodium pyruvate, 5% penicillin and streptomycin and 0% FBS) and cells were treated (15 µM surfen, 150 µM Condroitin Sulfate (CS) and 50 µM Dermatan Sulfate (DS)) for 48 hours. Photos were taken at 0 hours, 24 hours and 48 hours using a Floid Cell Imaging Station (Thermo Fisher Scientist), and cell migration was measured manually at different points of the wound. For collagen contraction we used a 48-well plate in which 30,000 cells were plated with 1mg/mL collagen normalized with PBS and NaOH. Once the mix was gelified, complete DMEM was added with different treatments (15 µM surfen, 150 µM Condroitin Sulfate (CS) and 50 µM Dermatan Sulfate (DS)). Photos were taken at different times and hours using a Floid Cell Imaging Station (Thermo Fisher Scientist) in order to observe the collagen contraction, that was measured manually.

### Chromatin immunoprecipitation (ChIP)

Fibroblasts were grown and chemically crosslinked by adding fresh 1% formaldehyde solution for 10 min at RT and quenched by adding 1/20 volume of 2.5 M of glycine for 5 min at RT. After washing twice in ice-cold PBS, cells were scraped and collected by centrifugation. The lysis of the cell pellet was in the lysis buffer (50 mM Tris-HCl (Ph 8), 10 mM EDTA, 1%SDS) for 30 min at 4°C. After lysis, cells were sonicated for 2X15 minutes (30-second ON, 30-second OFF pulses) at maximum power in a Bioruptor® Sonication System (Diagenode), and then centrifuged for high speed for 10 min. The supernatant containing the chromatin fraction was then subjected to immunoprecipitation or stored as input. Samples for immunoprecipitation were diluted 10 times using dilution buffer (0.01% SDS, 1.1% Triton X-100, 16.7 mM TRIS-HCl (pH8), 1.2 Mm EDTA). A blocking step was perfomed with protein G sepharose beads (1 hour) and after centrifugation, protein A/G beads together with the antibody or negative igG control were incubated with the samples overnight at 4°C. The following day, after different washes: Low Salt Buffer (0.1 %SDS, 0.1% TRITON X-100, 150mM NaCl, 2mM EDTA, 20mM TRIS-HCl (ph8)), High Salt Buffer (0.1 %SDS, 0.1% TRITON X-100, 500mM NaCl, 2mM EDTA, 20mM TRIS-HCl (ph8)) and Buffer TE (2X), the protein-DNA complexes were eluted from the beads twice by incubation with 125 μL of elution buffer (0.1M NaHCO3, 1% SDS) for 15 min with shaking. Reverse crosslinking was performed by adding a final concentration of 0.3M NaCl in the eluate and incubating it at 65°C during 4 hours with shaking. Thereafter, protein was removed from the samples by treating with proteinase K and 5X PK Buffer (50mM TRIS-HCl (pH8), 25mM EDTA, 1.25% SDS). DNA was subsequently purified by CHiP DNA clean & concentrator (ZYMO RESEARCH, #D5205) kit. In the case of input samples, they were also treated for crosslink reversal and the following steps.

### ChIP-seq analysis

Library construction and paired-end read sequencing (20M per sample) of triplicate immunoprecipitated chromatin (Cux1) and control inputs were performed using Illumina technology at STAB Vida (Portugal). After quality control, reads were aligned to human genome (hg38) using BWA. These BAM files were used for peak detection using MACS2, using pooled inputs and IPs and a FDR<=0.05. These regions were annotated using the chIPseekeR package in R, while HOMER software as used for motif detection.

### ReChIP

In the case of reChIP, the process was the same as in conventional chIP but using 3X of the number of cells. After washes the protein-DNA complexes were eluted from the beads twice by incubation with 50 μL of DTT 10mM for 10 min at 37°C with shaking. Samples were then diluted 10 times using dilution buffer (0.01% SDS, 1.1% Triton X-100, 16.7 mM TRIS-HCl (pH8), 1.2 Mm EDTA).

Samples again with the beads together with the antibody (Smad2/3 (Cell Signalling, #D7G7) or GADPH (Cell Signalling, #D16H11)) or igG negative control were incubated with the samples overnight at 4°C. Next day, the process followed as in conventional chIP.

### In silico docking

The molecular structures of human procathepsin L (PDB code: 1CS8) and chondroitin-4-sulfate (PDB code: 1C4S) were retrieved from Protein Databank. Docking was performed using Dockthor server with unbiased parameters. Visualization of the molecules was done using Pymol.

### Statistical analysis, availability of data and visualization

GraphPad Prism 8.0.2 and MATLAB software were used for representing and analysing data. Models were made using Biorender. Two-tailed Student’s T-test was performed for parametric variables, calculating P-value (P) to compare control conditions with various experimental groups. In Adhesion studies, paired two-tailed Student’s T-test was performed. The “n” value mentioned in the figure legends of quantitative polymerase chain reaction (qPCR) represents the number of independent samples. Asterisks mean: *P < 0.05; ***P < 0.01; ***P < 0.001; ****P < 0.0001. Raw and processed data for RNA-seq, chIP-seq and the rest of data can be found at 10.6084/m9.figshare.23185166.

## REFERENCES

Arthur, R. K., An, N., Khan, S., & McNerney, M. E. (2017). The haploinsufficient tumor suppressor, CUX1, acts as an analog transcriptional regulator that controls target genes through distal enhancers that loop to target promoters. Nucleic Acids Research, 45(11), 6350. https://doi.org/10.1093/NAR/GKX218

Bratulic, S., Limeta, A., Maccari, F., Galeotti, F., Volpi, N., Levin, M., Nielsen, J., & Gatto, F. (2022). Analysis of normal levels of free glycosaminoglycans in urine and plasma in adults. The Journal of Biological Chemistry, 298(2), 101575. https://doi.org/10.1016/J.JBC.2022.101575

Brokowska, J., Gaffke, L., Pierzynowska, K., Cyske, Z., & Węgrzyn, G. (2022). Cell cycle disturbances in mucopolysaccharidoses: Transcriptomic and experimental studies on cellular models. Experimental Biology and Medicine (Maywood, N.J.), 247(18), 1639–1649. https://doi.org/10.1177/15353702221114872

Carlson, M. A., & Longaker, M. T. (2004). The fibroblast-populated collagen matrix as a model of wound healing: a review of the evidence. Wound Repair and Regeneration, 12(2), 134–147. https://doi.org/10.1111/J.1067-1927.2004.012208.X

Chang, H. Y., Sneddon, J. B., Alizadeh, A. A., Sood, R., West, R. B., Montgomery, K., Chi, J. T., Van De Rijn, M., Botstein, D., & Brown, P. O. (2004). Gene Expression Signature of Fibroblast Serum Response Predicts Human Cancer Progression: Similarities between Tumors and Wounds. PLoS Biology, 2(2). https://doi.org/10.1371/JOURNAL.PBIO.0020007

Chatterjee, S., Stephenson, T. N., Michalak, A. L., Godula, K., & Huang, M. L. (2019). Silencing glycosaminoglycan functions in mouse embryonic stem cells with small molecule antagonists. Methods in Enzymology, 626, 249–270. https://doi.org/10.1016/bs.mie.2019.06.023

Costa, R., Urbani, A., Salvalaio, M., Bellesso, S., Cieri, D., Zancan, I., Filocamo, M., Bonaldo, P., Szabo, I., Tomanin, R., & Moro, E. (2017). Perturbations in cell signaling elicit early cardiac defects in mucopolysaccharidosis type II. Human Molecular Genetics, 26(9), 1643–1655. https://doi.org/10.1093/HMG/DDX069

Cyske, Z., Gaffke, L., Pierzynowska, K., & Węgrzyn, G. (2022). Complex Changes in the Efficiency of the Expression of Many Genes in Monogenic Diseases, Mucopolysaccharidoses, May Arise from Significant Disturbances in the Levels of Factors Involved in the Gene Expression Regulation Processes. Genes, 13(4). https://doi.org/10.3390/GENES13040593

De Pasquale, V., & Pavone, L. M. (2019). Heparan sulfate proteoglycans: The sweet side of development turns sour in mucopolysaccharidoses. Biochimica et Biophysica Acta. Molecular Basis of Disease, 1865(11). https://doi.org/10.1016/J.BBADIS.2019.165539

Gaffke, L., Pierzynowska, K., Krzelowska, K., Piotrowska, E., & Węgrzyn, G. (2020). Changes in expressions of genes involved in the regulation of cellular processes in mucopolysaccharidoses as assessed by fibroblast culture-based transcriptomic analyses. Metabolic Brain Disease, 35(8), 1353–1360. https://doi.org/10.1007/S11011-020-00614-2

Gaffke, L., Pierzynowska, K., Podlacha, M., Hoinkis, D., Rintz, E., Brokowska, J., Cyske, Z., & Wegrzyn, G. (2020). Underestimated aspect of mucopolysaccharidosis pathogenesis: Global changes in cellular processes revealed by transcriptomic studies. International Journal of Molecular Sciences, 21(4). https://doi.org/10.3390/ijms21041204

Gaffke, L., Szczudło, Z., Podlacha, M., Cyske, Z., Rintz, E., Mantej, J., Krzelowska, K., Węgrzyn, G., & Pierzynowska, K. (2022). Impaired ion homeostasis as a possible associate factor in mucopolysaccharidosis pathogenesis: transcriptomic, cellular and animal studies. Metabolic Brain Disease, 37(2), 299–310. https://doi.org/10.1007/S11011-021-00892-4

Gatto, F., Volpi, N., Nilsson, H., Nookaew, I., Maruzzo, M., Roma, A., Johansson, M. E., Stierner, U., Lundstam, S., Basso, U., & Nielsen, J. (2016). Glycosaminoglycan Profiling in Patients’ Plasma and Urine Predicts the Occurrence of Metastatic Clear Cell Renal Cell Carcinoma. Cell Reports, 15(8), 1822–1836. https://doi.org/10.1016/J.CELREP.2016.04.056

Goulet, B., Baruch, A., Moon, N. S., Poirier, M., Sansregret, L. L., Erickson, A., Bogyo, M., & Nepveu, A. (2004). A cathepsin L isoform that is devoid of a signal peptide localizes to the nucleus in S phase and processes the CDP/Cux transcription factor. Molecular Cell, 14(2), 207–219. https://doi.org/10.1016/S1097-2765(04)00209-6

Granados-Montiel, J., Cruz-Lemini, M., Rangel-Escareño, C., Martinez-Nava, G., Landa-Solis, C., Gomez-Garcia, R., Lopez-Reyes, A., Espinosa-Gutierrez, A., & Ibarra, C. (2021). SERPINA9 and SERPINB2: Novel Cartilage Lineage Differentiation Markers of Human Mesenchymal Stem Cells with Kartogenin. Cartilage, 12(1), 102. https://doi.org/10.1177/1947603518809403

Hampe, C. S., Polgreen, L. E., Lund, T. C., & McIvor, R. S. (2020). Dysostosis Multiplex in Human Mucopolysaccharidosis Type 1 H and in Animal Models of the Disease. Pediatric Endocrinology Reviews: PER, 17(4), 317–326. https://doi.org/10.17458/PER.VOL17.2020.HPL.DYSOSTOSISMULTIPLEXHUMANANIMAL

Henderson, N. C., Rieder, F., & Wynn, T. A. (2020). Fibrosis: from mechanisms to medicines. Nature, 587(7835), 555–566. https://doi.org/10.1038/S41586-020-2938-9

Heppner, J. M., Zaucke, F., & Clarke, L. A. (2015). Extracellular matrix disruption is an early event in the pathogenesis of skeletal disease in mucopolysaccharidosis I. Molecular Genetics and Metabolism, 114(2), 146–155. https://doi.org/10.1016/J.YMGME.2014.09.012

Huang, M. L., Michalak, A. L., Fisher, C. J., Christy, M., Smith, R. A. A., & Godula, K. (2018). Small Molecule Antagonist of Cell Surface Glycosaminoglycans Restricts Mouse Embryonic Stem Cells in a Pluripotent State. *Stem Cells (Dayton*, Ohio), 36(1), 45–54. https://doi.org/10.1002/STEM.2714

Iozzo, R. V, & Schaefer, L. (2015). Proteoglycan form and function: A comprehensive nomenclature of proteoglycans. Matrix Biology: Journal of the International Society for Matrix Biology, 42, 11–55. https://doi.org/10.1016/j.matbio.2015.02.003

Iyer, V. R., Eisen, M. B., Ross, D. T., Schuler, G., Moore, T., Lee, J. C. F., Trent, J. M., Staudt, L. M., Hudson, J., Boguski, M. S., Lashkari, D., Shalon, D., Botstein, D., & Brown, P. O. (1999). The transcriptional program in the response of human fibroblasts to serum. Science (New York, N.Y.), 283(5398), 83–87. https://doi.org/10.1126/SCIENCE.283.5398.83

Jiang, Z., Byers, S., Casal, M. L., & Smith, L. J. (2020). Failures of Endochondral Ossification in the Mucopolysaccharidoses. Current Osteoporosis Reports, 18(6), 759–773. https://doi.org/10.1007/S11914-020-00626-Y

Junqueira, L. C. U., & Montes, G. S. (1983). Biology of collagen-proteoglycan interaction. Archivum Histologicum Japonicum = Nihon Soshikigaku Kiroku, 46(5), 589–629. https://doi.org/10.1679/AOHC.46.589

Keller, C., Briner, J., Schneider, J., Spycher, M., Rampini, S., & Gitzelmann, R. (1987). [Mucopolysaccharidosis 6-A (Maroteaux-Lamy disease): comparison of clinical and pathologico-anatomic findings in a 27-year-old patient]. Helvetica Paediatrica Acta, 42(4), 317–333.

Kim, D.-W., & Lassar, A. B. (2003). Smad-Dependent Recruitment of a Histone Deacetylase/Sin3A Complex Modulates the Bone Morphogenetic Protein-Dependent Transcriptional Repressor Activity of Nkx3.2. Molecular and Cellular Biology, 23(23), 8704. https://doi.org/10.1128/MCB.23.23.8704-8717.2003

Klagsbrun, M., & Baird, A. (1991). A dual receptor system is required for basic fibroblast growth factor activity. Cell, 67(2), 229–231. https://doi.org/10.1016/0092-8674(91)90173-V

Lamberg, S. I., & Stoolmiller, A. C. (1974). Glycosaminoglycans. A Biochemical and Clinical Review. Journal of Investigative Dermatology, 63(6), 433–449. https://doi.org/10.1111/1523-1747.EP12680346

LeBleu, V. S., & Neilson, E. G. (2020). Origin and functional heterogeneity of fibroblasts. FASEB Journal, 34(3), 3519–3536. https://doi.org/10.1096/FJ.201903188R

Li, S. De, Moy, L., Pittman, N., Shue, G., Aufiero, B., Neufeld, E. J., LeLeiko, N. S., & Walsh, M. J. (1999). Transcriptional repression of the cystic fibrosis transmembrane conductance regulator gene, mediated by CCAAT displacement protein/cut homolog, is associated with histone deacetylation. Journal of Biological Chemistry, 274(12), 7803–7815. https://doi.org/10.1074/jbc.274.12.7803

Liu, N., Sun, Q., Wan, L., Wang, X., Feng, Y., Luo, J., & Wu, H. (2020). CUX1, A Controversial Player in Tumor Development. Frontiers in Oncology, 10, 534066. https://doi.org/10.3389/FONC.2020.00738/BIBTEX

Lizarraga, G., Lichtler, A., Upholt, W. B., & Kosher, R. A. (2002). Studies on the Role of Cux1 in Regulation of the Onset of Joint Formation in the Developing Limb. https://doi.org/10.1006/dbio.2001.0559

Lurje, I., Gaisa, N. T., Weiskirchen, R., & Tacke, F. (2023). Mechanisms of organ fibrosis: Emerging concepts and implications for novel treatment strategies. Molecular Aspects of Medicine, 92, 101191. https://doi.org/10.1016/J.MAM.2023.101191

Maas, C., & de Maat, S. (2021). Therapeutic SERPINs: Improving on Nature. Frontiers in Cardiovascular Medicine, 8, 648349. https://doi.org/10.3389/FCVM.2021.648349/BIBTEX

Mason, R. W., & Massey, S. D. (1992). Surface activation of pro-cathepsin L. Biochemical and Biophysical Research Communications, 189(3), 1659–1666. https://doi.org/10.1016/0006-291X(92)90268-P

Muenzer, J. (2011). Overview of the mucopolysaccharidoses. Rheumatology, 50(suppl_5), v4–v12. https://doi.org/10.1093/RHEUMATOLOGY/KER394

Munakata, H., Takagaki, K., Majima, M., & Endo, M. (1999). Interaction between collagens and glycosaminoglycans investigated using a surface plasmon resonance biosensor. Glycobiology, 9(10), 1023–1027. https://doi.org/10.1093/GLYCOB/9.10.1023

Nassar, L. R., Barber, G. P., Benet-Pagès, A., Pagès, P., Casper, J., Clawson, H., Diekhans, M., Fischer, C., Navarro Gonzalez, J., Hinrichs, A. S., Lee, B. T., Lee, C. M., Muthuraman, P., Nguy, B., Pereira, T., Nejad, P., Perez, G., Raney, B. J., Schmelter, D.,… Kent, W. J. (2023). The UCSC Genome Browser database: 2023 update. Nucleic Acids Research, 51(D1), D1188–D1195. https://doi.org/10.1093/NAR/GKAC1072

Nepveu, A. (2001). Role of the multifunctional CDP/Cut/Cux homeodomain transcription factor in regulating differentiation, cell growth and development. Gene, 270(1–2), 1–15. https://doi.org/10.1016/S0378-1119(01)00485-1

Paganini, C., Costantini, R., Superti-Furga, A., & Rossi, A. (2019). Bone and connective tissue disorders caused by defects in glycosaminoglycan biosynthesis: a panoramic view. The FEBS Journal, 286(15), 3008–3032. https://doi.org/10.1111/FEBS.14984

Parfrey, N. A., & Hutchins, G. M. (1986). Hepatic fibrosis in the mucopolysaccharidoses. The American Journal of Medicine, 81(5), 825–829. https://doi.org/10.1016/0002-9343(86)90353-0

Sansregret, L., & Nepveu, A. (2008). The multiple roles of CUX1: insights from mouse models and cell-based assays. Gene, 412(1–2), 84–94. https://doi.org/10.1016/J.GENE.2008.01.017

Schuksz, M., Fuster, M. M., Brown, J. R., Crawford, B. E., Ditto, D. P., Lawrence, R., Glass, C. A., Wang, L., Tor, Y., & Esko, J. D. (2008). Surfen, a small molecule antagonist of heparan sulfate. Proceedings of the National Academy of Sciences of the United States of America, 105(35), 13075–13080. https://doi.org/10.1073/PNAS.0805862105

Sodhi, H., & Panitch, A. (2021). Glycosaminoglycans in Tissue Engineering: A Review. Biomolecules, 11(1), 1–22. https://doi.org/10.3390/BIOM11010029

Somoza, R. A., Welter, J. F., Correa, D., & Caplan, A. I. (2014). Chondrogenic Differentiation of Mesenchymal Stem Cells: Challenges and Unfulfilled Expectations. Tissue Engineering. Part B, Reviews, 20(6), 596. https://doi.org/10.1089/TEN.TEB.2013.0771

Subramanian, A., Tamayo, P., Mootha, V. K., Mukherjee, S., Ebert, B. L., Gillette, M. A., Paulovich, A., Pomeroy, S. L., Golub, T. R., Lander, E. S., & Mesirov, J. P. (2005). Gene set enrichment analysis: A knowledge-based approach for interpreting genome-wide expression profiles. Proceedings of the National Academy of Sciences of the United States of America, 102(43), 15545–15550. https://doi.org/10.1073/PNAS.0506580102/SUPPL_FILE/06580FIG7.JPG

Tabib, T., Huang, M., Morse, N., Papazoglou, A., Behera, R., Jia, M., Bulik, M., Monier, D. E., Benos, P. V., Chen, W., Domsic, R., & Lafyatis, R. (2021). Myofibroblast transcriptome indicates SFRP2hi fibroblast progenitors in systemic sclerosis skin. Nature Communications, 12(1). https://doi.org/10.1038/S41467-021-24607-6

Tomasek, J. J., Gabbiani, G., Hinz, B., Chaponnier, C., & Brown, R. A. (2002). Myofibroblasts and mechano-regulation of connective tissue remodelling. Nature Reviews. Molecular Cell Biology, 3(5), 349–363. https://doi.org/10.1038/NRM809

Vadnais, C., Awan, A. A., Harada, R., Clermont, P. L., Leduy, L., Bérubé, G., & Nepveu, A. (2013). Long-range transcriptional regulation by the p110 CUX1 homeodomain protein on the ENCODE array. BMC Genomics, 14(1). https://doi.org/10.1186/1471-2164-14-258

Vallet, S. D., Berthollier, C., & Ricard-Blum, S. (2022). The glycosaminoglycan interactome 2.0. American Journal of Physiology - Cell Physiology, 322(6), C1271–C1278. https://doi.org/10.1152/AJPCELL.00095.2022/ASSET/IMAGES/LARGE/AJPCE LL.00095.2022_F005.JPEG

Varki, A., Cummings, R., Esko, J., Freeze, H., Hart, G., & Marth, J. (1999). Proteoglycans and Glycosaminoglycans. https://www.ncbi.nlm.nih.gov/books/NBK20693/

Wiśniewska, K., Gaffke, L., Krzelowska, K., Węgrzyn, G., & Pierzynowska, K. (2022). Differences in gene expression patterns, revealed by RNA-seq analysis, between various Sanfilippo and Morquio disease subtypes. Gene, 812. https://doi.org/10.1016/J.GENE.2021.146090

Wraith, J. E. (1995). The mucopolysaccharidoses: a clinical review and guide to management. Archives of Disease in Childhood, 72(3), 263. https://doi.org/10.1136/ADC.72.3.263

Zhang, Q., Wang, P., Fang, X., Lin, F., Fang, J., & Xiong, C. (2022). Collagen gel contraction assays: From modelling wound healing to quantifying cellular interactions with three-dimensional extracellular matrices. European Journal of Cell Biology, 101(3), 151253. https://doi.org/10.1016/J.EJCB.2022.151253

