## Supplemental figures for "Chondroitin and dermatan sulfate exposure induces a wound healing state in fibroblasts through Cux1-mediated SerpinB2 transcriptional repression"

### SUPPLEMENTARY FIGURE 1

```
                  1                                                    50
hARSB_reference   MGPRGAASLP RGPGPRRLLL PVVLPLLLLL LLAPPGSGAG ASRPPHLVFL
hARSB_patient     MGPRGAASLP RGPGPRRLLL PVVLPLLLLL LLAPPGSGAG ASRPPHLVFL

                  51                                                  100
hARSB_reference   LADDLGWNDV GFHGSRIRTP HLDALAAGGV LLDNYYTQPL CTPSRSQLLT
hARSB_patient     LADDLGWNDV GFHGSRIRTP HLDALAAGGV LLDNYYTQPL CTPSRSQLLT

                  101                                                 150
hARSB_reference   GRYQIRTGLQ HQIIWPCQPS CVPLDEKLLP QLLKEAGYTT HMVGKWHLGM
hARSB_patient     GRYQIRTGLQ HQIIWPCQPS CVPLDEKLLP QLLKEAGYTT HMVRKWHLGM

                  151                                                 200
hARSB_reference   YRKECLPTRR GFDTYFGYLL GSEDYYSHER CTLIDALNVT RCALDFRDGE
hARSB_patient     YRKECLPTRR GFDTYFGYLL GSEDYYSHER CTLIDALNVT RCALDFRDGE

                  201                                                 250
hARSB_reference   EVATGYKNMY STNIFTKRAI ALITNHPPEK PLFLYLALQS VHEPLQVPEE
hARSB_patient     EVATGYKNMY STNIFTKRAI ALITNHPPEK PLFLYLALQS VHEPLQVPEE

                  251                                                 300
hARSB_reference   YLKPYDFIQD KNRHHYAGMV SLMDEAVGNV TAALKSSGLW NNTVFIFSTD
hARSB_patient     YLKPYDFIQD KNRHHYAGMV SLMDEAVGNV TAALKSSGLW NNTVFIFSTD

                  301                                                 350
hARSB_reference   NGGQTLAGGN NWPLRGRKWS LWEGGVRGVG FVASPLLKQK GVKNRELIHI
hARSB_patient     NGGQTLAGGN NWPLRGRKWS LWEGGVRGVG FVASPLLKQK GVKNRELIHI

                  351                                                 400
hARSB_reference   SDWLPTLVKL ARGHTNGTKP LDGFDVWKTI SEGSPSPRIE LLHNIDPNFV
hARSB_patient     SDWLPTLVKL ARGHTNGTKP LDGFDVWKTI SEGSPSPRIE LLHNIDPNFV

                  401                                                 450
hARSB_reference   DSSPCPRNSM APAKDDSSLP EYSAFNTSVH AAIRHGNWKL LTGYPGCGYW
hARSB_patient     DSSPCPRNSM APAKDDSSLP EYSAFNTSVH AAIRHGNWKL LTGYPGCGYW

                  451                                                 500
hARSB_reference   FPPPSQYNVS EIPSSDPPTK TLWLFDIDRD PEERHDLSRE YPHIVTKLLS
hARSB_patient     FPPPSQYNVS EIPSSDPPTK TLWLFDIDRD PEERHDLSRE YPHIVTKLLS

                  501                              533
hARSB_reference   RLQFYHKHSV PVYFPAQDPR CDPKATGVWG PWM
hARSB_patient     RLQFYHKHSV PVYFPAQDPR CDPKATGVWG PWM
```

Alignment of the reference human Arsb protein and the sequence with the homozygous mutation that is present in the ML patient (G144R).

**SUPPLEMENTARY FIGURE 2**

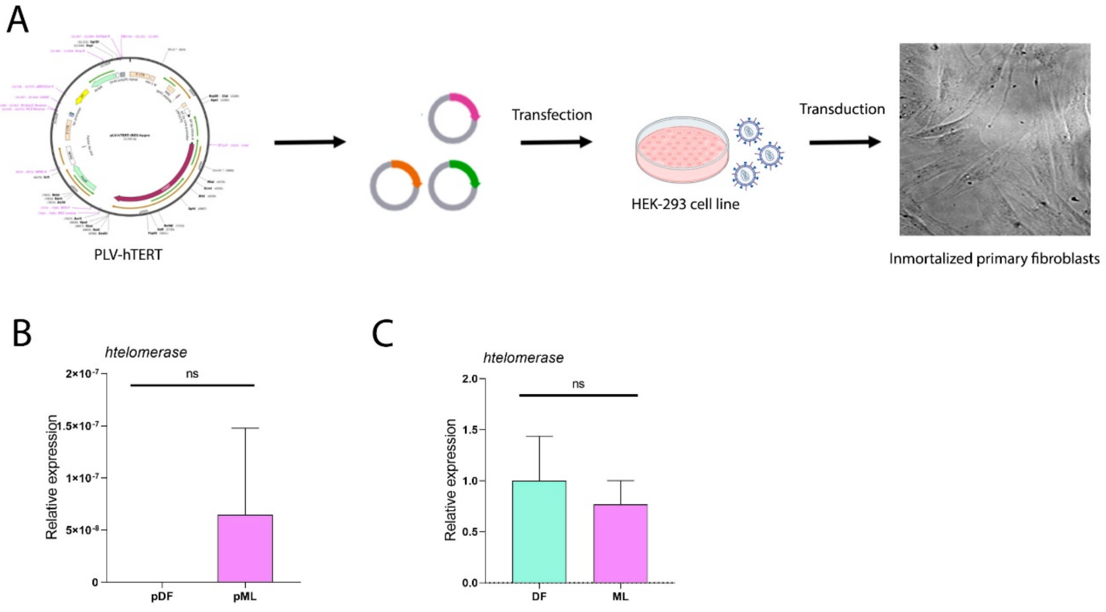


(A) Workflow of the immortalization process that was developed to generate DF and ML lines. (B-C) mRNA expression levels in DF vs ML. hTERT in primary cells (B), hTERT in immortalized cells as raw expression compared to constitutive gene (C). Average expression normalized to DF and standard deviation as error bars were plotted, n=6. P values were obtained using two-tailed t-test (ns means no significative differences).

**SUPPLEMENTARY FIGURE 3**

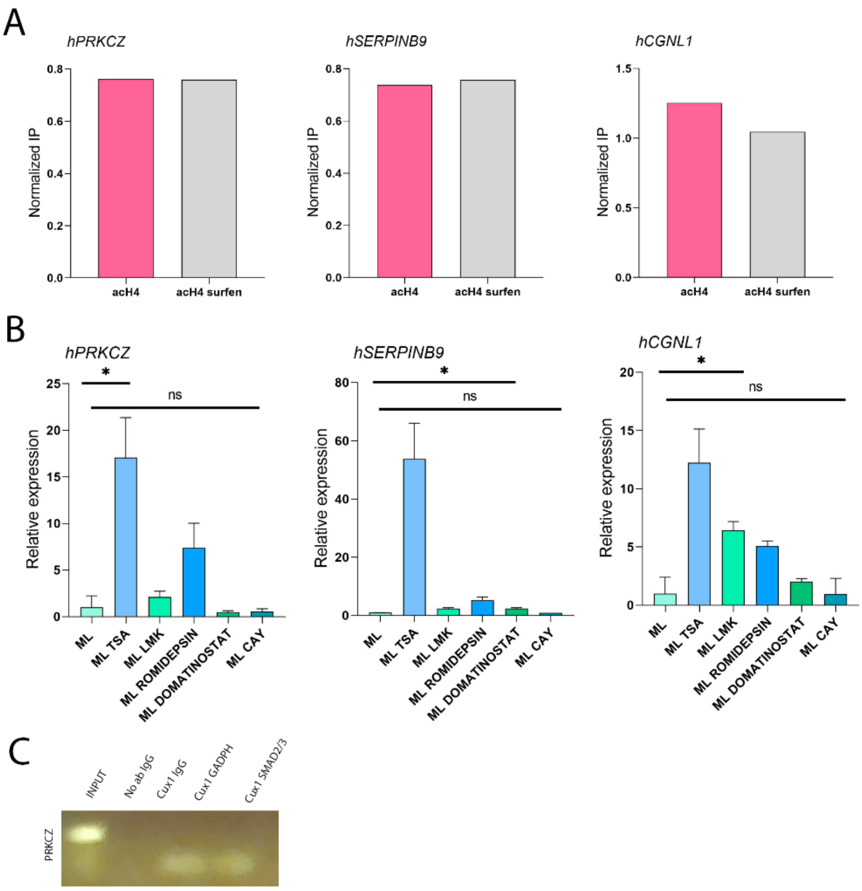


(A) Quantification of Acetyl H4 levels in ML cells with and without surfen treatment from chIP in hPRKCZ, hSERPINB9 and hCGNL1 regions. (B) mRNA expression levels in ML with different Histone Deacetylase (HDAC) inhibitors treatments hPRKCZ, hSERPINB9 and hCGNL1. Average expression is relative to untreated ML cells and standard deviation as error bars were plotted, n=3. (C) reChIP using Cux1 and GADPH or Cux1 and Smad2/3 antibodies in the PRKCZ region (Input and igG controls are included). P-values were obtained using two-tailed t-test (* represents p<0,05 and ns means no significative differences).

**SUPPLEMENTARY FIGURE 4**

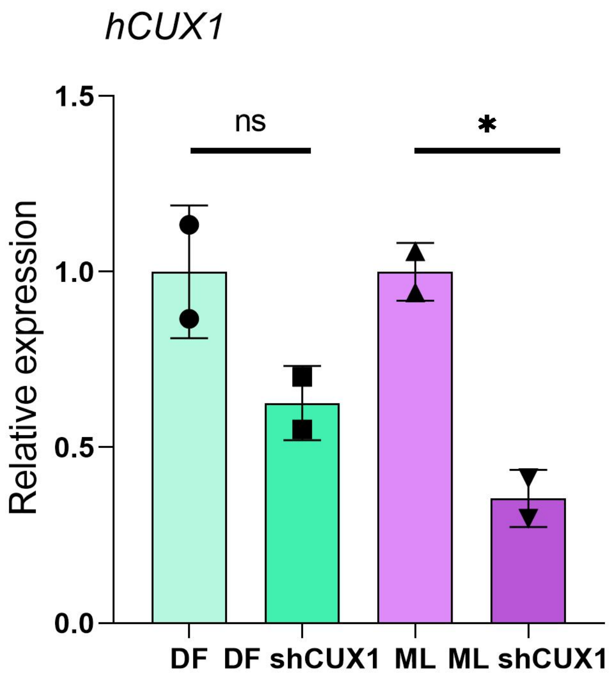


mRNA expression levels of CUX1 in DF and ML infected with the shCUX1 lentivirus, after 72 hours treated with doxycyclin. Average expression is relative to control cells and standard deviation as error bars were plotted, n=3.

**SUPPLEMENTARY FIGURE 5**

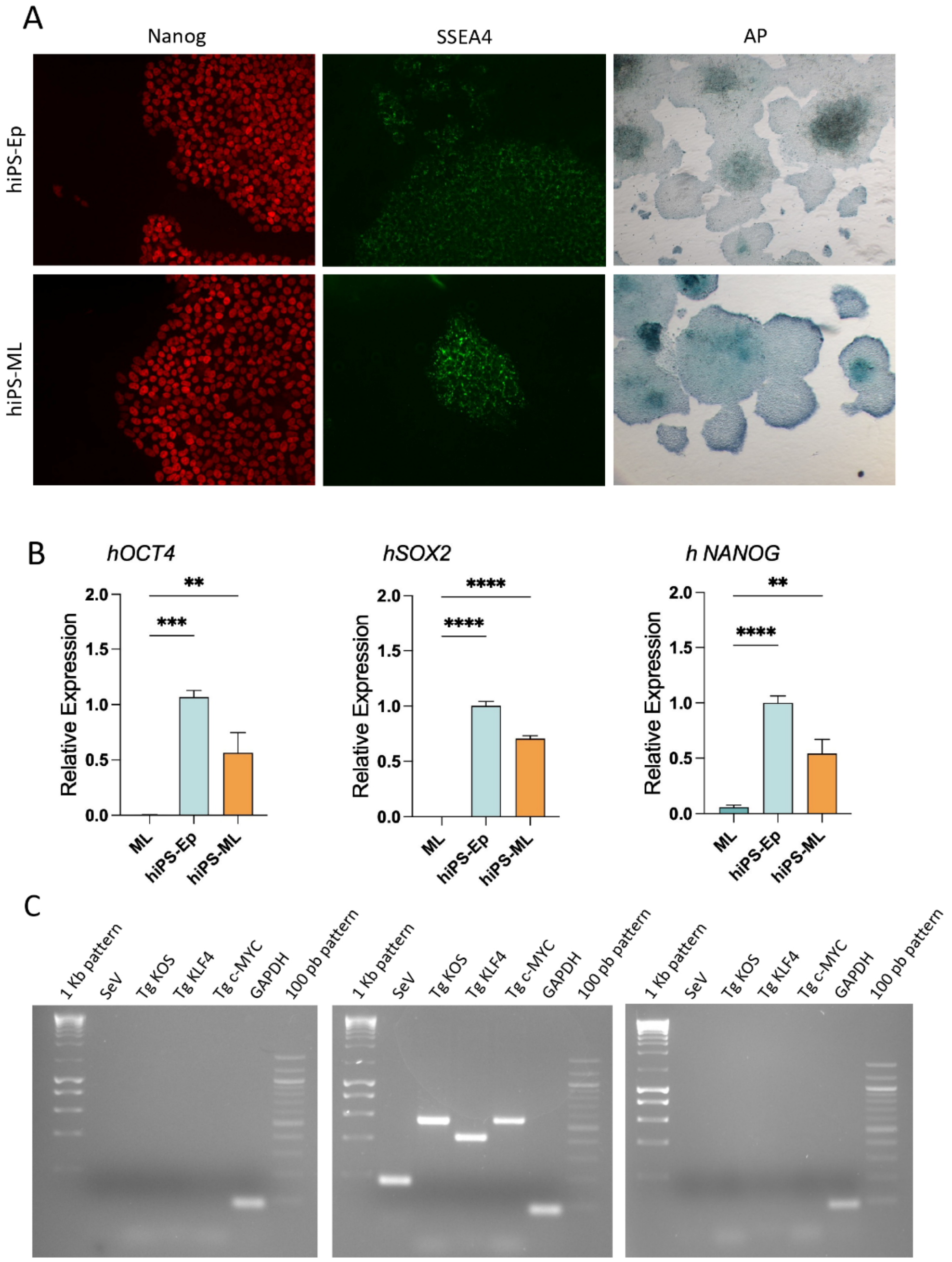


(A) Immunofluorescence from hiPS-ML for TFs NANOG and SSEA4. A' Characterization of the pluripotency of the generated iPSC line analyzing the activity of the enzyme alkaline phosphatase (AP). (B) mRNA expression levels in OCT4, SOX2 and NANOG. Mean expression is relative to episomal iPS cells and standard deviation as error bars were plotted, n=3. P-values were obtained using two-tailed t-test (**** represents p<0,0001, *** represents p<0,001, ** represents p<0,01). (C) RT-PCR analysis showing silencing of the transgenes KLF4, c-MYC, OCT4 and SOX2 and the absence of Sendai virus in the hiPS-ML line (right panel) compared to negative (left) and positive (center) controls.
